# Influence of neck tissue conductivities on the phrenic nerve activation threshold during non-invasive electrical stimulation

**DOI:** 10.1101/2025.05.12.653397

**Authors:** Laureen Wegert, Luca Di Rienzo, Lorenzo Codecasa, Marek Ziolkowski, Alexander Hunold, Tim Kalla, Irene Lange, Jens Haueisen

## Abstract

Phrenic nerve stimulation can be used as an artificial ventilation method to reduce the adverse effects of mechanical ventilation. Detailed computational models and electromagnetic simulations are used to determine appropriate stimulation parameters. Therefore, tissue parameters have to be selected, but they vary widely in the literature.

Here, we evaluated the phrenic nerve activation threshold using minimum and maximum electrical conductivity values found in the literature of each modeled neck tissue type. To calculate the phrenic nerve activation threshold, an anatomical detailed finite element model of the neck and a biophysiological nerve model were used.

Considerable changes in nerve activation thresholds were found for the following tissue conductivities (with decreasing effects): muscle, skin, soft tissue, subcutaneous fat, and nerve tissue. Changes in the nerve activation threshold due to changes in skin conductivity occurred due to the bridging effect, which is an unwanted and avoidable effect during stimulation.

In conclusion, fat, muscle, nerve, and soft tissue require the most accurate tissue properties and geometric representation within the model.

## 1. Introduction

Artificial ventilation is a life-saving technique commonly used in intensive care units. Current methods of mechanical ventilation stress the lungs and can cause atrophy of the main inspiratory muscle, the diaphragm. Electrical stimulation of the phrenic nerve, which innervates the diaphragm, keeps the muscle active and results in physiological respiratory movements. Non-invasive stimulation of the phrenic nerve can be performed electrically in the neck region [1].

Choosing the optimal stimulation parameters is challenging and electromagnetic simulations are used to select appropriate parameters [2]. Therefore, detailed models and accurate tissue parameters are required.

However, biological tissue properties are highly dependent on intra- and inter-individual differences, as well as the measurement conditions such as the temperature and the frequency [3]. For this reason, a wide range of permittivity and conductivity values is found in the literature. The use of different tissue properties strongly influences the results of the electromagnetic field calculation and leads to different potential distributions and nerve activation thresholds [4–6].

Here, we investigated the influence of neck tissue conductivities on the phrenic nerve activation threshold during non-invasive electrical stimulation. Therefore, we evaluated the resulting nerve activation threshold when sweeping from the minimum to the maximum conductivity values of each tissue type.

## 2 Materials & Methods

We developed a complex, anatomical detailed volume conductor model of the neck, comprising 12 tissue types — skin, subcutaneous fat, soft tissue, muscle, bone, cartilage, intervertebral disc, blood, thyroid, trachea, esophagus, and nerve — spanning from the clavicle to the jaw, and bounded by cut planes. The model, along with the stimulation electrodes [2] and the tissue compartments, is depicted in Figure 1.

**Fig. 1.**
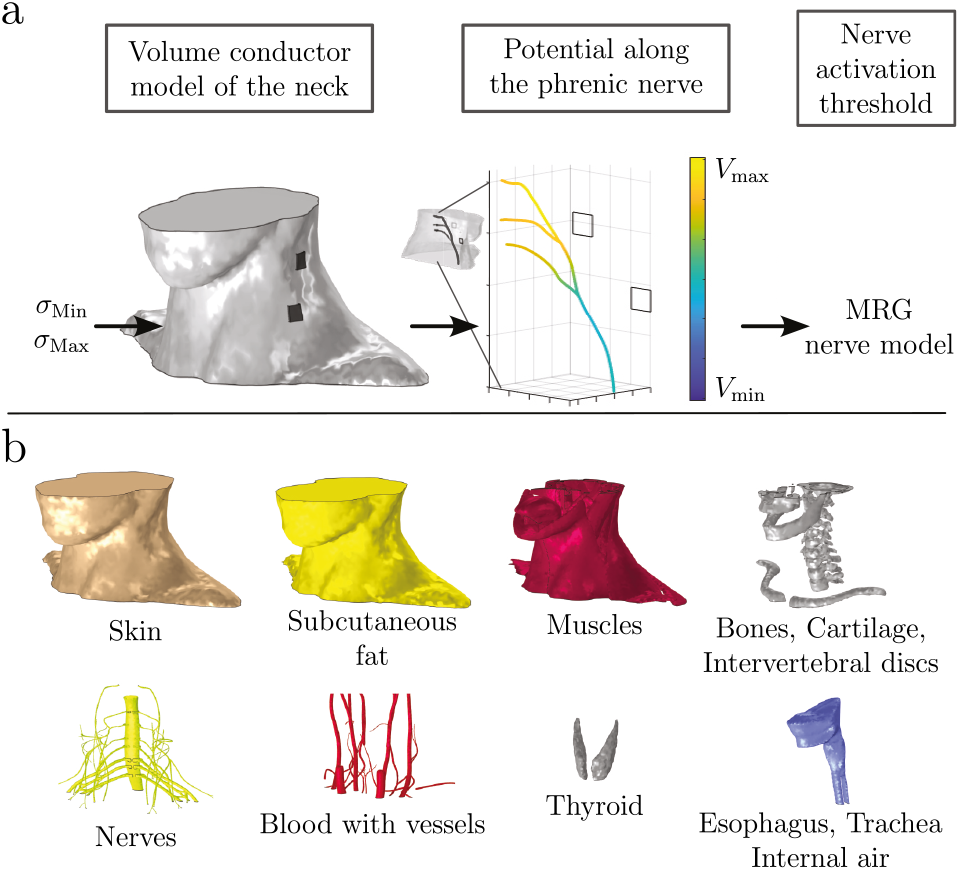
a) Overview of the model pipeline used to calculate the phrenic nerve activation threshold. The volume conductor model of the neck was used to calculate the potential distribution along the phrenic nerve with minimum and maximum conductivities of each tissue type, followed by calculation of the nerve activation threshold via the McIntyre-Richardson-Grill (MRG) nerve model. Electrodes were marked with squares. Upper electrode was used as anode, bottom electrode as cathode. b) Tissue types used within the model were skin, subcutaneous fat, muscle, bone, cartilage, intervertebral disc, nerve (incl. phrenic nerve), blood, thyroid, esophagus, trachea, and internal air. Space between the compartments was filled with soft tissue.

The stimulation current was applied via two surface electrodes (size 1 cm × 1 cm, center-center distance 3.3 cm) positioned at the posterior border of the sternocleidomastoid muscle. At the stimulation electrodes, a normal current density of 200 A*/*m^2^ was applied. The ground potential was set at an additional electrode at the back of the neck. To ensure potential current flow through the cut planes into the head and the thorax, floating potentials were assigned at each cut plane. The mesh consisted of approximately 94 million tetrahedral elements. Material properties were assigned for each tissue according to the mean, minimum, and maximum conductivities from Table 1. The calculation of electric potential was performed under steady-state conditions using the Electric Currents interface of COMSOL Multiphysics (V. 6.0.405, COMSOL AB, Stockholm, Sweden). The electric potential along the phrenic nerve was extracted along the midlines of the fascicles reaching the roots C3, C4, and C5. For each fascicle, the nerve activation threshold was calculated with the McIntyre-Richardson-Grill (MRG) nerve model for an illustrative stimulation signal (cathodic, monophasic, pulse width 150 µs) and an sample axon fiber diameter of 10 µm in the NEURON simulation environment (V. 8.2.2).

**Table 1.** Conductivity values for the tissue types used within the model. Minimum and maximum conductivities were chosen from [7–9]. Mean conductivities were from a previous study [2].

| Tissue | Mean (mS/m) | Min (mS/m) | Max (mS/m) |
| --- | --- | --- | --- |
| Skin | 0.251 | 0.2 | 250 |
| Fat | 41.28 | 24.0 | 215 |
| Soft Tissue | 342.8 | 79.2 | 389 |
| Muscle | 298.6 | 100.0 | 726 |
| Bone | 20.3 | 1.85 | 20.8 |
| Cartilage | 172.4 | 161 | 1430 |
| Intervertebral Disc | 830 | 600 | 1430 |
| Nerve (incl. Phrenic Nerve) | 39.42 | 17.1 | 852 |
| Thyroid | 524.8 | 481 | 537 |
| Esophagus | 480.8 | 164 | 536 |
| Trachea | 310 | 300 | 342 |
| Blood | 700.6 | 433 | 946 |
| Electrodes | 3.33 | 0.1 | 1000 |

To investigate the influence of uncertain tissue conductivity on the nerve activation threshold, the minimum and maximum conductivity values were derived from the literature [7–9] and summarized in Table 1. For each tissue, the conductivity was iteratively varied to the minimum and to the maximum value. For the other tissues, the mean conductivity value from a previous study was used [2]. The potential along the phrenic nerve and the mean phrenic nerve activation threshold for the three fascicles were calculated for each set of conductivity values. The differences in the nerve activation threshold using the minimum and the maximum conductivity value of each tissue were evaluated and compared to the reference nerve activation threshold based on mean tissue conductivity value.

For tissues with changes in the nerve activation threshold of *>* 1 mA, ten additional values per decade were sampled equidistantly between the minimum and maximum conductivity to examine the relationship between conductivity and nerve activation threshold.

## 3 Results

Varying conductivities resulted in changes in the nerve activation threshold from the reference value of 37 mA calculated with the mean conductivities. The minimum and maximum values of the nerve activation threshold values corresponding to the conductivity changes are presented in Figure 2.

**Fig. 2.**
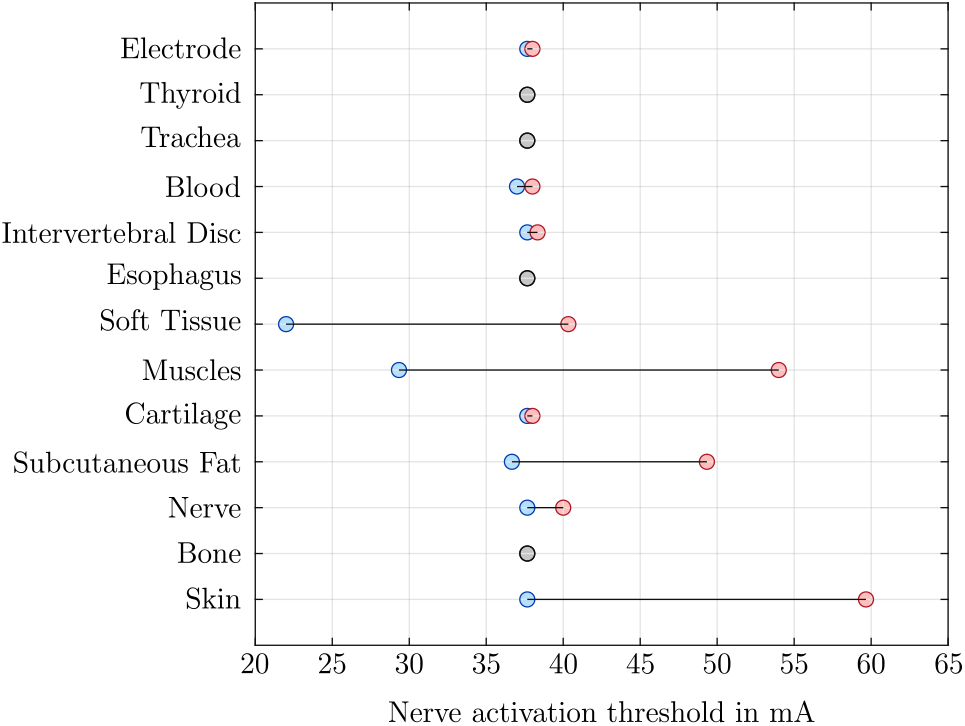
Nerve activation threshold for each tissue with minimum and maximum conductivity. For the tissue types soft tissue, muscle, nerve, and skin, variations higher than 1 mA compared to the reference nerve activation threshold with mean conductivities (37 mA) were observed.

No considerable changes from the reference activation threshold were observed for the tissues thyroid, trachea, esophagus, and bone. Small changes of *≤* 1 mA were observed for the tissues blood, intervertebral disc, cartilage, and the electrode. Conductivity changes in nerve tissue resulted in a changed nerve activation threshold of 3 mA for maximum conductivity. The largest changes were 25 mA, 22 mA, 18 mA, and 12 mA for muscle, skin, soft tissue, and subcutaneous fat.

Using the minimum conductivity resulted in a decreased nerve activation threshold (marked in blue) and using the maximum conductivity resulted in an increased threshold (marked in red). The lowest nerve activation threshold was observed at the minimum conductivity of soft tissue with 22 mA, the highest nerve activation threshold for maximum conductivity of skin tissue with 59 mA.

For the tissues skin, subcutaneous fat, soft tissue, muscle, and nerve tissue, additional sample points between the minimum and maximum conductivity were calculated. The results are shown in Figure 3 with a logarithmic conductivity axis.

**Fig. 3.**
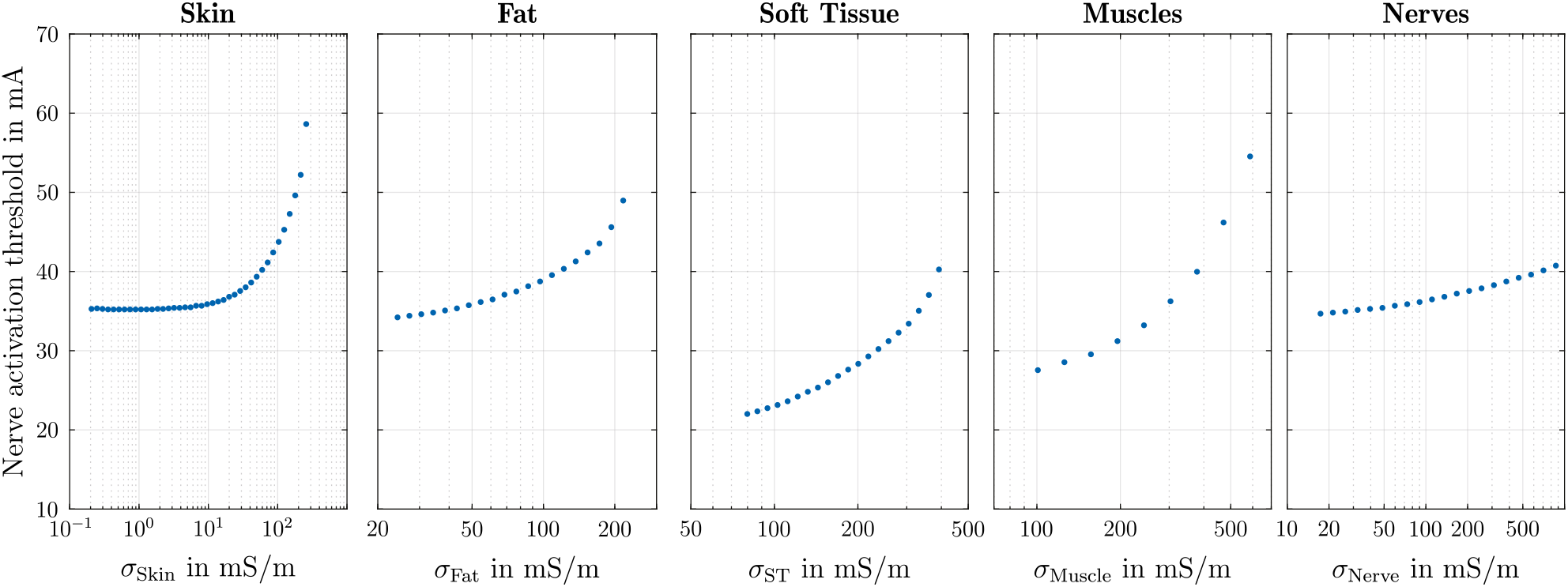
Nerve activation thresholds from sampled conductivity values between minimum and maximum conductivity. Increasing the conductivity resulted in higher nerve activation threshold for all tissue types. For skin tissue, increased nerve activation threshold started at conductivities higher than 10 mS*/*m.

Nerve activation thresholds ranged from 37 mA to 59 mA for changes in skin conductivity. A constant nerve activation threshold was observed from minimum conductivity up to approximately 10 mS*/*m. For changes in fat tissue conductivity, the nerve activation threshold increased steadily from 37 mA to 49 mA. Similarly, for soft tissue conductivity, the threshold increased from 22 mA to 40 mA with a steady rise. Increasing muscle conductivity resulted in nerve activation thresholds between 29 mA and 54 mA, with a consistent increment. For nerve conductivity, the smallest change was observed, increasing from 37 mA to 40 mA. Across all five tissues, a smooth transfer function was observed.

## 4 Discussion

The neck model’s dominant tissues — skin, fat, soft tissue, muscle, and nerve — exhibited changes in nerve activation thresholds, consistent with literature [4, 5]. Their proximity to both the current injection electrodes and the phrenic nerve evaluation path explains their substantial impact on potential distribution and consequently on the nerve activation threshold, a phenomenon also documented in electroencephalography studies [4, 6].

As reported by Klein et al. [5], utilizing the minimum conductivity of the surrounding tissue led to a decrease in the nerve activation threshold.

Changes in nerve activation thresholds due to skin conductivity variations were noticeable only when skin conductivity was close to its maximum. Under these conditions, the skin’s higher conductivity relative to the underlying subcutaneous fat tissue created a current bridge, directing the majority of the current between electrodes. This resulted in an increased current requirement for phrenic nerve activation, consistent with the loss effect reported in the review by Keller and Kuhn about experimental and modeling studies [9]. Similar threshold changes would be observed when conductive solutions or gels were applied to the entire skin, indicating this should be avoided.

The model presented herein is limited by its single variable conductivity analysis, precluding the investigation of interactive effects among tissue conductivities. A detailed investigation of conductivity uncertainty and its impact on nerve activation thresholds would require further Monte Carlo analysis.

## Conclusion

To access the impact of altered tissue conductivity, minimum and maximum values were used. Significant changes in nerve activation thresholds were observed across muscle, skin, subcutaneous fat, soft tissue, and nerve.

Accurate conductivity measurements of influential tissues and precise geometric modeling are essential, as individual tissue conductivities can lead to distinct nerve activation thresholds.

The observed steady relationship between conductivity and nerve activation thresholds suggests the applicability of a generalized polynomial chaos model [6], which will be explored in future work.

## Author Statement

### Research funding

The author received funding from the Federal Ministry of Education and Research with the grant *eVent* (13GW0591B).

